# A Comprehensive Model of Glucose-Insulin Regulation Including Acute and Prolonged Effects of Physical Activity in Type 1 Diabetes

**DOI:** 10.1101/2021.06.09.447693

**Authors:** Julia Deichmann, Sara Bachmann, Marc Pfister, Gabor Szinnai, Hans-Michael Kaltenbach

## Abstract

**Objective:** For type 1 diabetic patients, accurate adjustment of insulin treatment to physical activity (PA) is a challenging open problem. Glucose uptake by the exercising muscles increases acutely, causing increased hepatic glucose production to maintain glucose homeostasis. Meanwhile, insulin sensitivity is elevated for a prolonged period to drive glycogen repletion during recovery. These processes strongly depend on PA duration and intensity, making their combined effects difficult to predict accurately. In this work, we develop a model of glucose-insulin regulation that captures PA from low to high intensity including acute and prolonged effects on glucose metabolism.

**Methods:** We extended an existing minimal model of glucose-insulin regulation to capture PA-driven changes in glucose metabolism. We incorporated the insulin-independent increase in glucose uptake and production, including the effects of glycogen depletion and of high-intensity PA on production. The model also captures the prolonged increase in insulin sensitivity.

**Results:** The model accurately predicts glucose dynamics of published data during short and prolonged PA of moderate to high intensity and during subsequent recovery. In-silico full-day studies elucidate the effects of timing, duration and intensity of PA and of insulin bolus reduction on glucose levels during and after the activity.

**Conclusion:** The proposed model captures the blood glucose dynamics during all main PA regimes.

**Significance:** Mathematical models of glucose-insulin regulation are critical components of closed-loop insulin delivery and clinical decision support systems for achieving good glycemic control. The presented model shows potential for the development and assessment of algorithms targeting treatment adjustment to PA.

## 1 Introduction

Type 1 diabetes (T1D) results from autoimmune destruction of pancreatic *β*-cells. Affected patients depend on lifelong treatment by exogenous insulin to achieve good glycemic control.

Physical activity (PA) is beneficial for T1D patients and recommended in clinical guidelines [5,14], but fear of acute and late-onset (often nocturnal) hypoglycemia restrains many patients from exercising [7]. Preventing hypoglycemia by adjusting insulin treatment and nutrition is a major challenge due to the complexity of PA-driven changes in glucose metabolism, and current guidelines consider only coarse categories of glycemia, PA duration and intensity to determine the recommended adjustments [1, 13, 37].

Exercise-induced processes occur on different time-scales and strongly depend on duration and intensity of PA [39]. During PA, the glucose demand by active muscles increases acutely and glucose uptake (GU) from plasma is up-regulated. Simultaneously, hepatic glucose production (GP) by gluconeogenesis and glycogenolysis increases to maintain plasma glucose levels [9, 26]. During prolonged PA, liver glycogen stores may deplete and GP cannot be maintained, causing a drop in glucose levels [19]. In contrast, GP may (initially) exceed GU and result in rising plasma glucose during high-intensity PA due to an increase in catecholamines and cortisol [32]. During recovery, insulin-independent GU and GP rates quickly return to their baseline levels [9]. However, insulin sensitivity can stay elevated for up to 48 hours after PA to replete liver glycogen stores and may cause late-onset hypoglycemia, including nocturnal hypoglycemia [33].

Mathematical models allow us to determine clinical parameters such as insulin sensitivity from an intravenous glucose tolerance test [6] and to better understand the glucose-insulin system in physiological detail [41]. They also play a critical role in the development of decision support systems and of closed-loop insulin delivery systems (artificial pancreas) [11, 28, 29].

The need to include the impact of PA on glucose metabolism in diabetes-related models has long been recognized [13,38,42], but the prolonged and nonlinear effects during exercise and recovery pose formidable challenges. Existing models are usually restricted to moderate intensity and often focus exclusively on either short-term or prolonged effects of PA. To-date, no model simultaneously incorporates all acute and prolonged changes in glucose metabolism caused by exercise from low to high intensity, which limits the applicability to predict exercise-induced hypoglycemia and to determine accurate treatment adjustments.

A first extension of the Bergman minimal model [6] includes constant intensity exercise with acute increases in glucose uptake and production and eventual liver glycogen depletion, but not the elevated insulin sensitivity during recovery [40]. An alternative extension by Breton allows different exercise intensities for the insulin-independent increase in glucose clearance but not for insulin sensitivity, and does not consider glycogen depletion [8]. Starting from their earlier glucose-insulin model with carbohydrate intake, Dalla Man and coworkers integrate Breton’s exercise model and add intensity-dependent insulin sensitivity [15, 16].

Two physiologically detailed models with exercise include glucose uptake and production, but no changes in insulin sensitivity [30], respectively glycogen depletion with increased hepatic glucose uptake after a meal to drive glycogen repletion after PA [21]. Based on the latter, a virtual patient population generator capturing PA has been proposed [36]. Furthermore, multi-scale models of tissue and organ systems connected by the circulatory system have been considered [27, 35].

Here, we propose a semi-mechanistic model of glucose-insulin regulation that captures the acute and prolonged changes in glucose metabolism during PA and subsequent recovery for low-to high-intensity exercise. We use a well-established two-compartment minimal model [12] as our core model for glucose-insulin dynamics and combine it with simple models of insulin kinetics [34] and carbohydrate absorption. We then incorporate the separate acute increase in glucose uptake and production—including depletion—and the prolonged rise in insulin sensitivity caused by PA. We use sigmoidal transfer functions to compactly describe some of the physiological changes. Our model accurately predicts blood glucose dynamics during and after PA and could be used to evaluate the associated hypoglycemia risk several hours post-exercise. Its modularity allows easy extensions of the insulin and meal submodels, which makes the developed model suitable for incorporation into both decision support systems (‘exercise calculator’ [13]) with bolus injections and pump-based artificial pancreas systems.

## 2 Methods

Our model consists of a core model including basic glucose-insulin regulation, insulin bolus kinetics, and meal absorption, and an exercise model capturing the PA-related changes in glucose metabolism.

### 2.1 Core Model

#### 2.1.1 Two-compartment minimal model

The two-compartment minimal model [12] describes the impact of insulin and glucose on plasma glucose levels with compartment *Q*_1_ [mg/kg] to represent glucose mass in plasma, and a remote compartment *Q*_2_ [mg/kg] (Fig. 1). Plasma insulin promotes the disappearance of plasma glucose into liver and tissue, and suppresses hepatic glucose production via the dynamic state *X* [1/min]:

**Figure 1:**
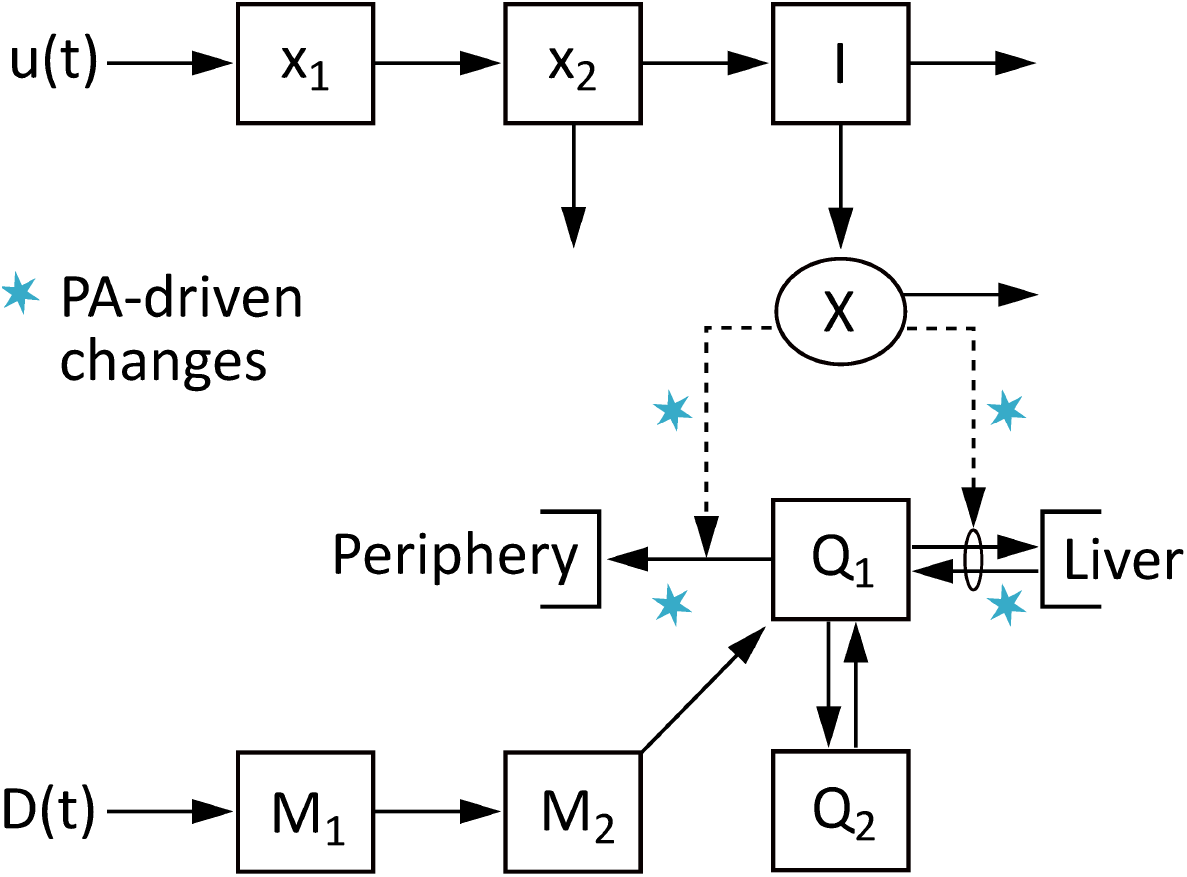
Schematic of the glucose-insulin model used to describe glucose (*Q*_1_) and insulin (*I*) dynamics. It is based on the two-compartment minimal model [12], extended to capture plasma insulin kinetics [34] after injection *u* and to capture the glucose response to a meal *D. X* is a dynamic state related to insulin concentration. Processes affected by PA are marked with a blue asterisk.

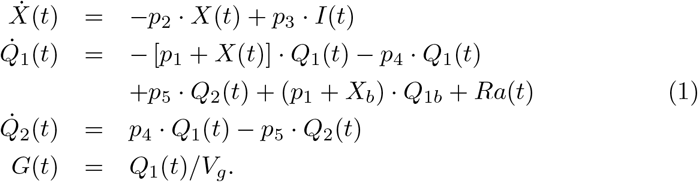

Here, *I* [*µ*U/ml] is the plasma insulin concentration, *Q*_1*b*_ [mg/kg] denotes the basal level of glucose and *X*_*b*_ the basal value of *X* with *X*_*b*_ = *p*_3_*/p*_2_ *·I*_*b*_, *Ra* [mg/kg/min] is the rate of glucose appearance in plasma after a meal. *G* [mg/dl] is the plasma glucose concentration, *V*_*g*_ [dl/kg] the glucose distribution volume and *p*_*i*_ are rate parameters.

#### 2.1.2 Insulin kinetics model

We use a model with two subcutaneous compartments of insulin masses *x*_1_ and *x*_2_ [*µ*U] and a plasma insulin compartment *I*_*c*_ [*µ*U/ml] (Fig. 1) to capture the resulting plasma insulin kinetics after a subcutaneous bolus injection [34]:

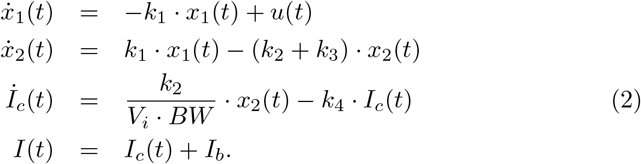

Insulin is injected into *x*_1_ at rate *u* [*µ*U/min] and yields a rise *I*_*c*_ in plasma insulin concentration *I*, adjusted for distribution volume *V*_*i*_ [ml/kg] and body-weight *BW* [kg]. *k*_*i*_ are rate parameters and we assume a constant basal insulin level *I*_*b*_ [*µ*U/ml].

#### 2.1.3 Meal absorption model

We describe glucose appearance after a meal with a two-compartment absorption model:

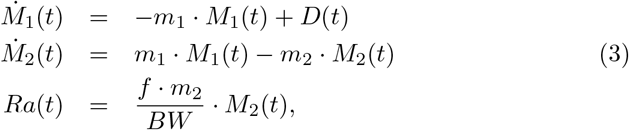

where the ingested glucose, *D* [mg/min], first passes through compartments *M*_1_ [mg] and *M*_2_ [mg] before being absorbed (Fig. 1). *Ra* [mg/kg/min] defines the rate of glucose appearance in plasma as the fraction *f* of absorbed glucose; *m*_*i*_ are rate parameters.

### 2.2 Exercise Model

To incorporate exercise-driven changes, we replace the plasma glucose mass equation (1) with

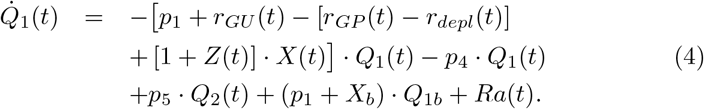

The rates *r*_*GU*_ and (*r*_*GP*_ *−r*_*depl*_) provide the insulin-independent increase in GU and GP (including glycogen depletion), respectively, while (1+*Z*) captures the PA-driven increase in insulin sensitivity.

We model smooth transitions between different exercise modes, e.g. low- and high-intensity PA, via the transfer function

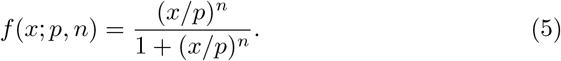

Although this transfer function does not offer a physiologically accurate description of the underlying molecular mechanisms, it enables us to keep the model simple, while incorporating the effects of different exercise regimes on glucose levels. This strategy has been used before to activate the exercise-driven increase in insulin sensitivity [8].

#### 2.2.1 Measure of exercise intensity and duration

We consider movement via accelerometer (AC) counts, which we link to PA intensity *Y* via a delay *τ*_*AC*_ [min] to allow initial adaptation to PA:

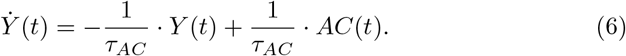

Furthermore, we track PA duration *t*_*PA*_ [min], integrated AC count *PA*_*int*_ [counts] and time spent at high intensity *t*_*HI*_ [min]:

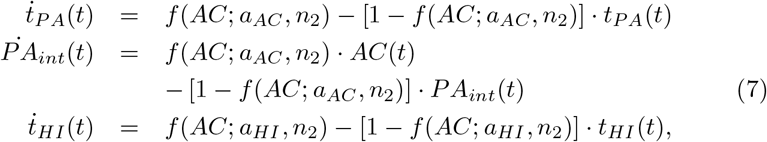

where transfer functions *f*(*AC*; *a*_*AC*_, *n*_2_) and *f*(*AC*; *a*_*HI*_, *n*_2_) capture the transition from rest to exercise, respectively from low to high intensity.

#### 2.2.2 Insulin sensitivity

We model the rise in insulin sensitivity as

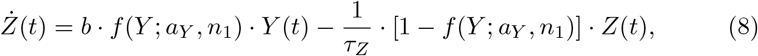

where *f*(*Y* ; *a*_*Y*_, *n*_1_) defines the minimal intensity *Y* considered as PA; *b* and *τ*_*Z*_ are parameters. Insulin sensitivity increases proportionally with intensity during PA, stays elevated to drive glycogen repletion after PA, and then returns slowly to its baseline.

#### 2.2.3 Insulin-dependent glucose uptake and production

The PA-driven, insulin-independent glucose uptake (*r*_*GU*_ [1/min]) and production (*r*_*GP*_ [1/min]) rates are given by

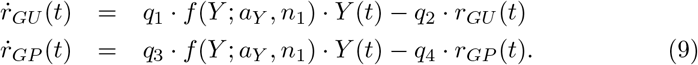

Both rates depend on the PA intensity and are used to model a quick-on, quick-off behavior with rate parameters *q*_*i*_. Parameters *q*_3_ and *q*_4_ are modulated between low-(subscript ‘LI’) and high-intensity (subscript ‘HI’) values by a transfer function to smoothly transition between the two exercise regimes:

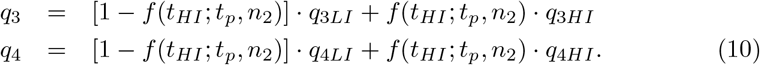

#### 2.2.4 Glycogen depletion

We determine the time *t*_*depl*_ [min] until depletion of liver glycogen stores from the integrated AC count and PA duration:

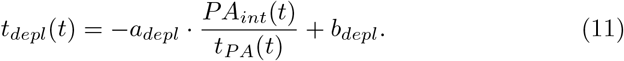

After depletion, GP decreases with rate *r*_*depl*_ [1/min]:

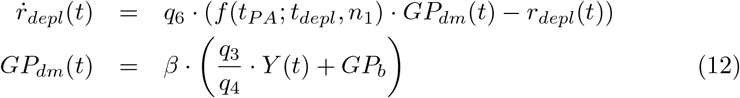

and the transfer function indicates whether exercise time exceeded *t*_*depl*_. The maximum drop *GP*_*dm*_ in GP is the sum of the PA-driven GP at steady state, *q*_3_*/q*_4_ · *Y* (*t*), and the basal GP rate, *GP*_*b*_. *q*_6_ is a rate parameter and *β* is the proportion of net hepatic glucose production attributed to glycogenolysis.

### 2.3 Parameter Determination

We obtained parameter values from literature or physiological knowledge when feasible. We estimated the remaining parameters from published data using least squares regression. Parameter values are given in Table 1.

**Table 1:**
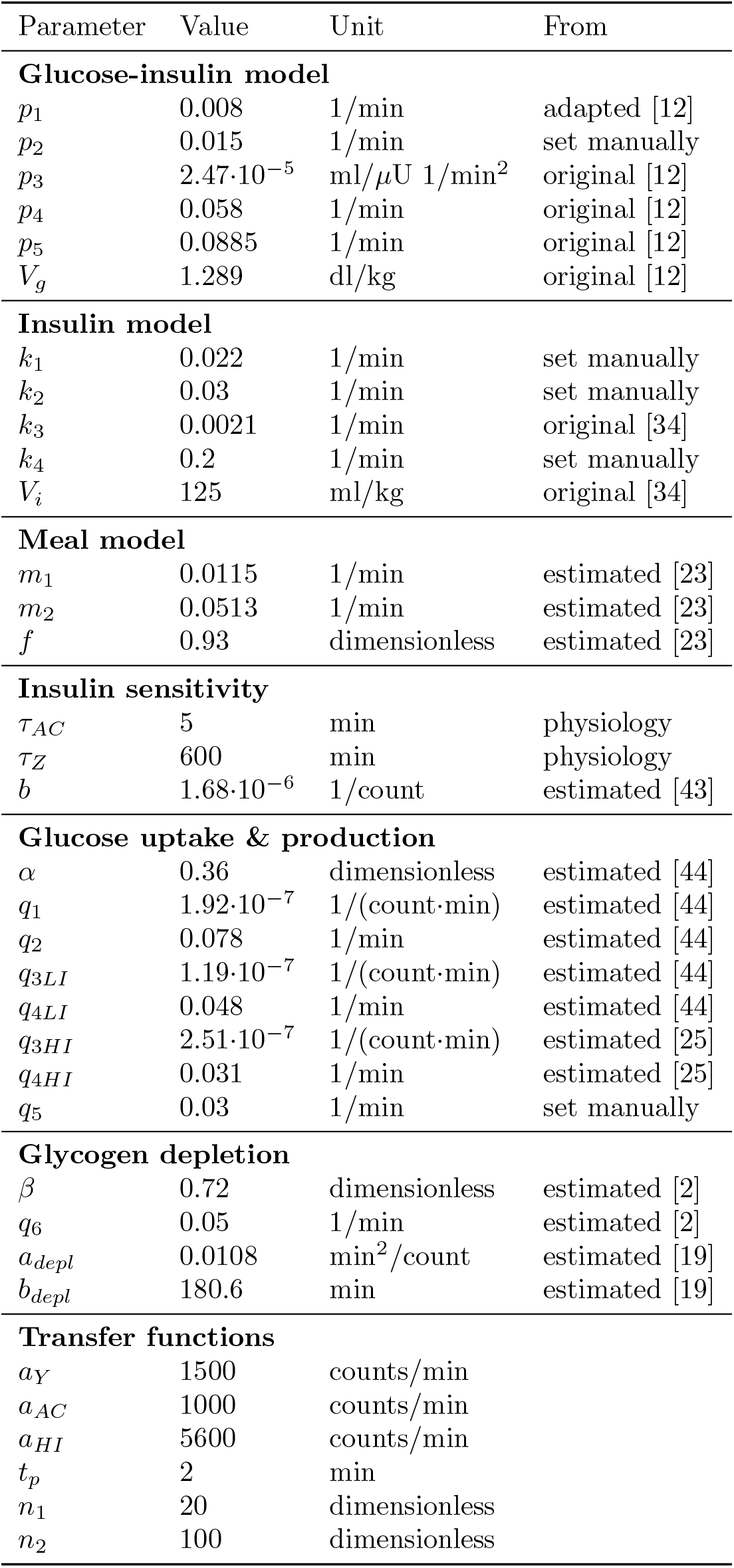
Model Parameters

| Parameter | Value | Unit | From |
| --- | --- | --- | --- |
| **Glucose-insulin model** |  |  |  |
| $p_1$ | 0.008 | 1/min | adapted [12] |
| $p_2$ | 0.015 | 1/min | set manually |
| $p_3$ | $2.47 \cdot 10^{-5}$ | ml/μU 1/min<sup>2</sup> | original [12] |
| $p_4$ | 0.058 | 1/min | original [12] |
| $p_5$ | 0.0885 | 1/min | original [12] |
| $V_g$ | 1.289 | dl/kg | original [12] |
| **Insulin model** |  |  |  |
| $k_1$ | 0.022 | 1/min | set manually |
| $k_2$ | 0.03 | 1/min | set manually |
| $k_3$ | 0.0021 | 1/min | original [34] |
| $k_4$ | 0.2 | 1/min | set manually |
| $V_i$ | 125 | ml/kg | original [34] |
| **Meal model** |  |  |  |
| $m_1$ | 0.0115 | 1/min | estimated [23] |
| $m_2$ | 0.0513 | 1/min | estimated [23] |
| $f$ | 0.93 | dimensionless | estimated [23] |
| **Insulin sensitivity** |  |  |  |
| $\tau_{AC}$ | 5 | min | physiology |
| $\tau_Z$ | 600 | min | physiology |
| $b$ | $1.68 \cdot 10^{-6}$ | 1/count | estimated [43] |
| **Glucose uptake & production** |  |  |  |
| $\alpha$ | 0.36 | dimensionless | estimated [44] |
| $q_1$ | $1.92 \cdot 10^{-7}$ | 1/(count·min) | estimated [44] |
| $q_2$ | 0.078 | 1/min | estimated [44] |
| $q_{3LI}$ | $1.19 \cdot 10^{-7}$ | 1/(count·min) | estimated [44] |
| $q_{4LI}$ | 0.048 | 1/min | estimated [44] |
| $q_{3HI}$ | $2.51 \cdot 10^{-7}$ | 1/(count·min) | estimated [25] |
| $q_{4HI}$ | 0.031 | 1/min | estimated [25] |
| $q_5$ | 0.03 | 1/min | set manually |
| **Glycogen depletion** |  |  |  |
| $\beta$ | 0.72 | dimensionless | estimated [2] |
| $q_6$ | 0.05 | 1/min | estimated [2] |
| $a_{depl}$ | 0.0108 | min<sup>2</sup>/count | estimated [19] |
| $b_{depl}$ | 180.6 | min | estimated [19] |
| **Transfer functions** |  |  |  |
| $a_Y$ | 1500 | counts/min | |
| $a_{AC}$ | 1000 | counts/min | |
| $a_{HI}$ | 5600 | counts/min | |
| $t_p$ | 2 | min | |
| $n_1$ | 20 | dimensionless | |
| $n_2$ | 100 | dimensionless | |

We are aware that already the simple two-compartment minimal model of glucose-insulin regulation is known to be unidentifiable [12], and our model is therefore unlikely to be identifiable from blood glucose measurements alone. Nevertheless, we found that the reported parameter values enable accurate prediction of plasma glucose levels for a wide variety of experimental scenarios (see below). To calibrate our model, we have to rely on literature values and domain knowledge for some parameters. However, we calibrated the model parts related to physical activity based on published data sets. We separated parameters into process-specific sets and individually estimated these on data sets acquired during the corresponding exercise modes.

We used experimental data from different published studies to test if our calibrated model is able to accurately describe independent data. Importantly, these studies not only provide independent data not used in calibration of our model, but also investigated exercise at different intensities and for varying durations compared to the data sets used for parameter estimation. This enables us to investigate the general applicability of our model to various PA types and provides further validation of the intensity dependence given in the model.

We also tested whether model predictions qualitatively agree with clinical knowledge over an extended period of time by performing several full-day simulations with different PA scenarios and with and without insulin bolus adjustments.

#### 2.3.1 Core model parameters

We slightly modified the original core model and manually adjusted insulin parameters to mimic the appearance of the plasma insulin peak after about 60 min, typical for current fast-acting insulins. We estimated *m*_1_, *m*_2_ and *f* of the meal model from plasma glucose measurements after a 75g oral glucose load [23].

#### 2.3.2 Exercise model parameters

We set the delay parameter *τ*_*AC*_ to 5 min [8] and chose a time constant *τ*_*Z*_ of 600 min such that insulin sensitivity stays elevated for up to 48h in accordance with literature reports [33].

We estimated the increase in insulin sensitivity during PA (parameter *b*) from measurements of the insulin-dependent rate of glucose disappearance during rest and 100 min of cycling at 80% VO_2_^max^ in healthy subjects [43]. We converted %VO_2_^max^ to accelerometer count using

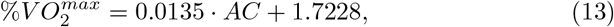

estimated from simultaneous AC count and %VO_2_^max^ measurements for different types and intensities of PA [17].

We estimated the insulin-independent GU and GP parameters *q*_1_, *q*_2_, *q*_3*LI*_ and *q*_4*LI*_ from total GU and GP rates measured in healthy adults during 60 min of PA at 40% VO_2_^max^ [44]. We distinguished between resting and exercise-driven contributions by separating the net rate of glucose change at rest into endogenous glucose production and glucose uptake:

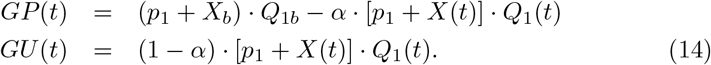

We estimated the high-intensity exercise parameters *q*_3*HI*_ and *q*_4*HI*_ from interstitial glucose measurements [25] of T1D patients performing 45 min of intervals at 82.5% VO_2_^max^ and introduced the parameter *q*_5_ = 0.03 min^-1^ to prevent a switch to low-intensity parameters during recovery.

We determined the time until hepatic glycogenolysis decreases due to glycogen depletion from reported depletion times for different intensities [19] (parameters *a*_*depl*_ and *b*_*depl*_). We estimated glycogen depletion parameters *β* and *q*_6_ from plasma glucose measurements [2] recorded during 3h of cycling at 58% VO_2_^max^ in healthy individuals, where we restricted *q*_6_ to 0.05 min^-1^ to avoid an overshoot in GP after PA.

Finally, we enforce the transition from rest to PA between 1000 and 2000 counts/min with parameters *a*_*Y*_ = 1500 counts/min and *n*_1_ = 20 [17]. Accordingly, we defined *a*_*AC*_ = 1000 counts/min and *n*_2_ = 100 to track duration and AC count immediately from the start of PA. High-intensity PA commences at 80% VO_2_^max^ (5800 counts/min), and we set *a*_*HI*_ = 5600 counts/min and *t*_*p*_ = 2 min for a transition between intensity regimes at 75%-80% VO_2_^max^.

## 3 Results

### 3.1 Glucose Metabolism during Moderate-Intensity Exercise

We first evaluated the effects of moderate-intensity PA on plasma glucose dynamics on a cycling session of 60 min at 40% VO_2_^max^, the data used for estimating the GU and GP rates [44]. Model and observed data are in very good agreement (Fig. 2).

**Figure 2:**
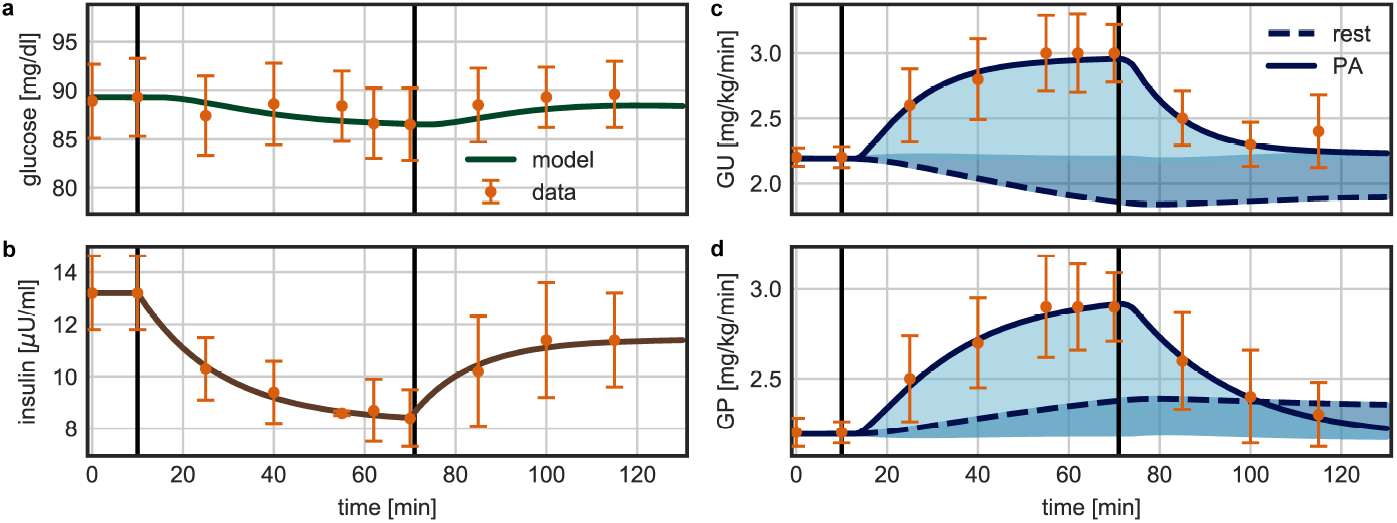
Data [44] and model prediction for 60 min of moderate-intensity PA (vertical lines) at 40% VO_2_^max^. (a) Model prediction of plasma glucose concentration. (b) Plasma insulin concentration. Model fit of (c) glucose uptake (GU) and (d) glucose production (GP) rates during PA. The difference to resting rates is separated into contributions of insulin-dependent (dark shaded area) and insulin-independent (bright shaded area) changes in glucose metabolism due to PA.

Both GU and GP rates increase at the onset of PA, starting from a baseline level of 2.2 mg/kg/min. GU increases to 3.0 mg/kg/min due to a higher uptake by the exercising muscles, while GP increases to 2.9 mg/kg/min to meet the increased demand. During recovery, both rates return quickly to their pre-exercise levels.

In our model, we separate the PA-driven changes in GU and GP into insulin-dependent and -independent contributions. While insulin-independent GU and GP rise immediately at the beginning of PA and turn off quickly after the end of exercise, insulin sensitivity increases gradually during the activity and stays elevated during recovery, leading to a continued rise in GU and drop in GP compared to resting rates. Plasma insulin concentration decreases during PA in the healthy individual to counteract the effects of increased insulin sensitivity (while T1D patients must actively reduce the insulin dose), and increases again after PA to drive glycogen repletion. Glucose homeostasis is maintained throughout the activity, where plasma glucose levels only decrease slightly from 89.3 mg/dl to 86.5 mg/dl, and return to previous levels after the activity.

We validated the calibrated exercise model on data from two additional independent studies in healthy subjects. In the first study [22], participants cycled for 60 min at 60% VO_2_^max^. Glucose levels decreased from 93.7 mg/dl to 86.5 mg/dl and remained constant during the first hour of recovery. In the second study [4], participants performed arm exercise on a cycle ergometer for 120 min at 30% VO_2_^max^ and glucose concentration remained stable throughout PA. We again observe good agreement between data and model predictions (Fig. 3).

**Figure 3:**
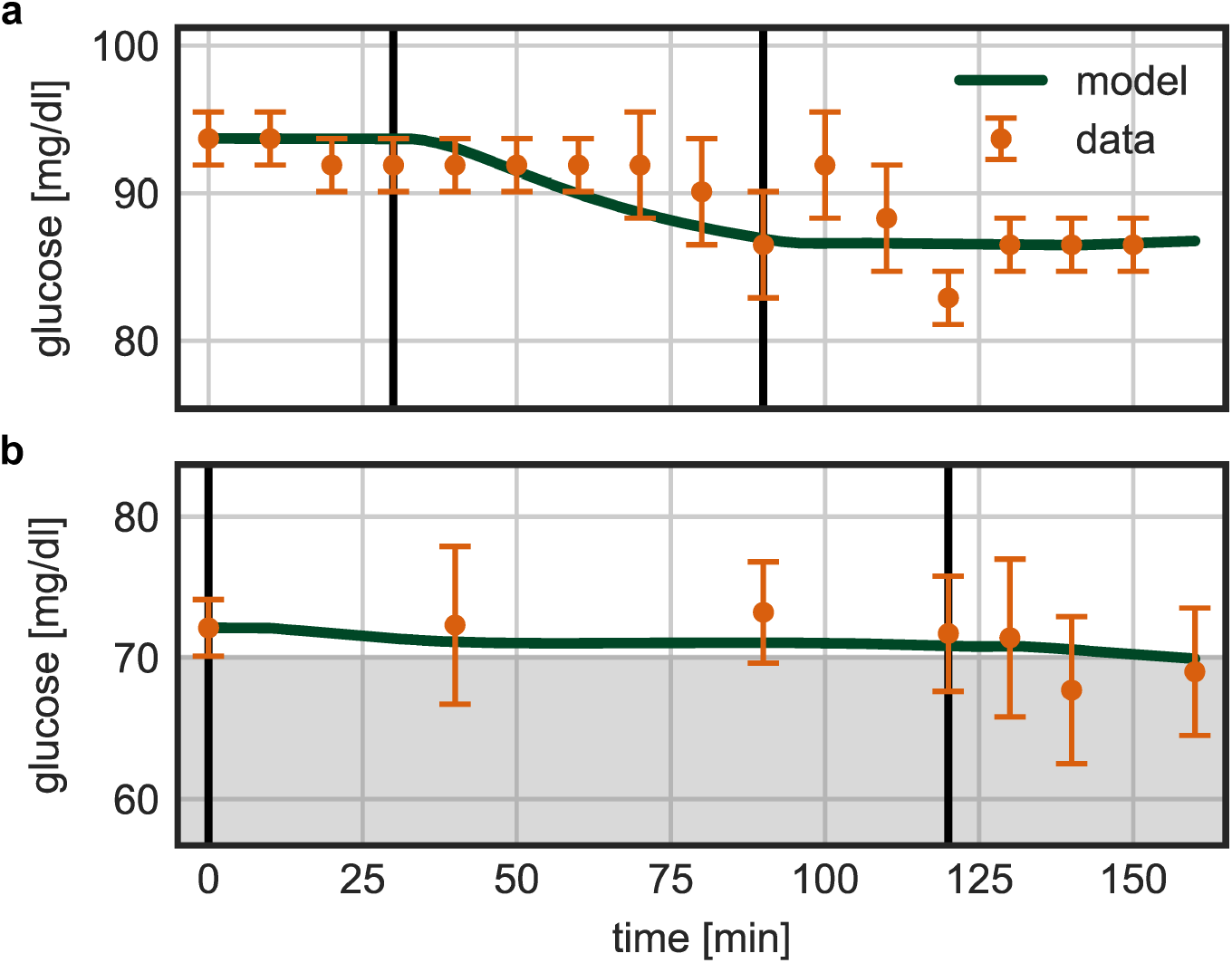
Validation of exercise model. (a) Model prediction and data [22] for 60 min of PA (vertical lines) at 60% VO_2_^max^. (b) Model prediction and data [4] for 120 min of PA at 30% VO_2_^max^.

### 3.2 Glycogen Depletion during Prolonged Exercise

To evaluate the dynamics of glycogen depletion, we compared model predictions to the data for 180 min of PA at 58% VO_2_^max^ previously used for parameter estimation [2]. During the first two hours of PA, plasma glucose concentration decreases slowly from 79.1 mg/dl to 64 mg/dl. In the last 60 min, GP drops below its pre-exercise levels due to depletion, causing glucose levels to drop quickly reaching 50.1 mg/dl at the end of PA. During recovery, GP returns to the baseline level and plasma glucose starts rising. The model correctly captures glycogen depletion after 136 min (Eq. 11) and the observed plasma glucose levels (Fig. 4a).

**Figure 4:**
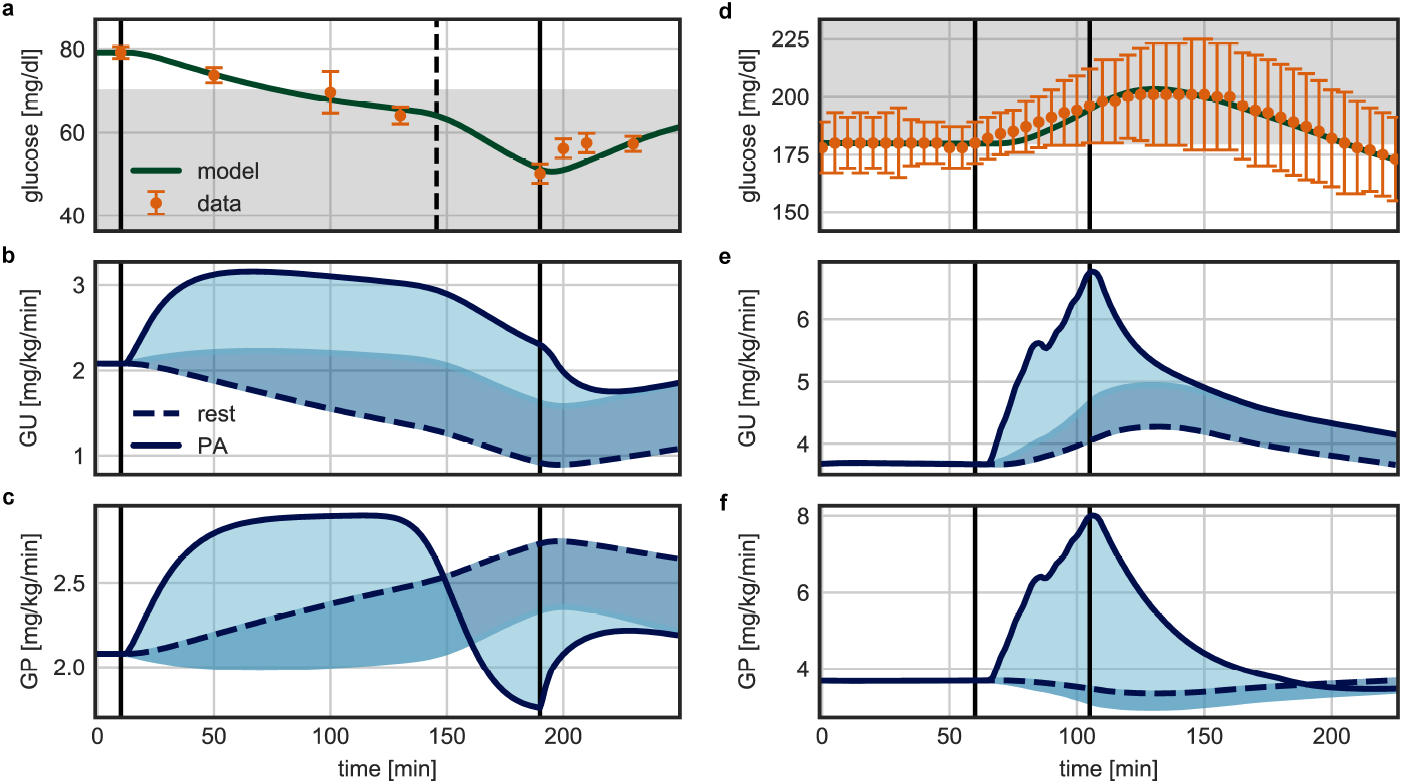
(a-c) Data [2] and model prediction for 180 min of PA (vertical lines) at 58% VO_2_^max^. (d-f) Data [25] and model prediction for 45 min of high-intensity PA at 82.5% VO_2_^max^. (a, d) Model fits of plasma glucose concentration, where the dashed vertical line indicates time of depletion. Model predictions of (b, e) GU and (c, f) GP during PA. The difference to resting rates is separated into contributions of insulin-dependent (dark shaded area) and insulin-independent (bright shaded area) changes in glucose metabolism due to PA.

Importantly, we find good agreement of our model predictions with the observed data for the time before onset of depletion, even though we did not use this part of the data for model calibration. We also determined the time of depletion independently and again observe good agreement for these data.

For independent validation of the depletion model, we used a data set of plasma glucose levels measured in healthy adults during 240 min of cycling at 30% VO_2_^max^ [3]. At this intensity, depletion is reached after 158 min. The model correctly predicts the drop in glucose levels from 81.3 mg/dl at the beginning to 56.2 mg/dl at the end of PA (Fig. 5).

**Figure 5:**
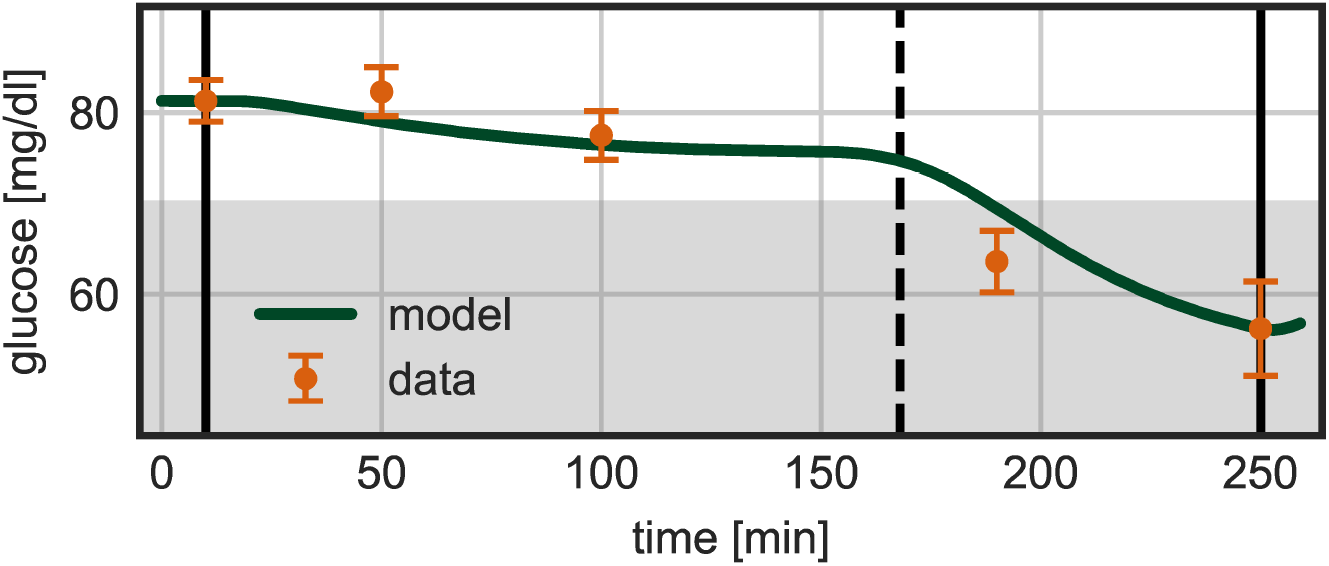
Model prediction and data [3] for 240 min of PA (vertical lines) at 30% VO_2_^max^ for validation of the exercise model. The dashed vertical line indicates time of depletion.

### 3.3 Glucose Dynamics during High-Intensity Exercise

We used the previous data of high-intensity intervals at 82.5% VO_2_^max^ for 45 min to evaluate this part of the exercise model [25]. The initial glucose level is 180 mg/dl and increases during PA to a maximum of 201 mg/dl shortly after PA before returning to pre-exercise levels. GP increases from 3.7 mg/kg/min at rest to 8.0 mg/kg/min during PA, while GU reaches 6.8 mg/kg/min. This dynamics is correctly reflected by the model (Fig. 4b).

### 3.4 Full-Day Simulation of Glucose Dynamics in T1D

Overall, our model adequately reflects experimental data for a range of exercise scenarios and generalizes to independent scenarios. We next considered its potential for long-term predictions for a full day of a T1D patient, consisting of three meals and corresponding insulin boluses. We defined twelve PA scenarios for this typical day: exercise in the morning or afternoon, of duration 60 or 180 min and with moderate (30% or 60%) or high (90% VO_2_^max^) intensity (Fig. 6).

**Figure 6:**
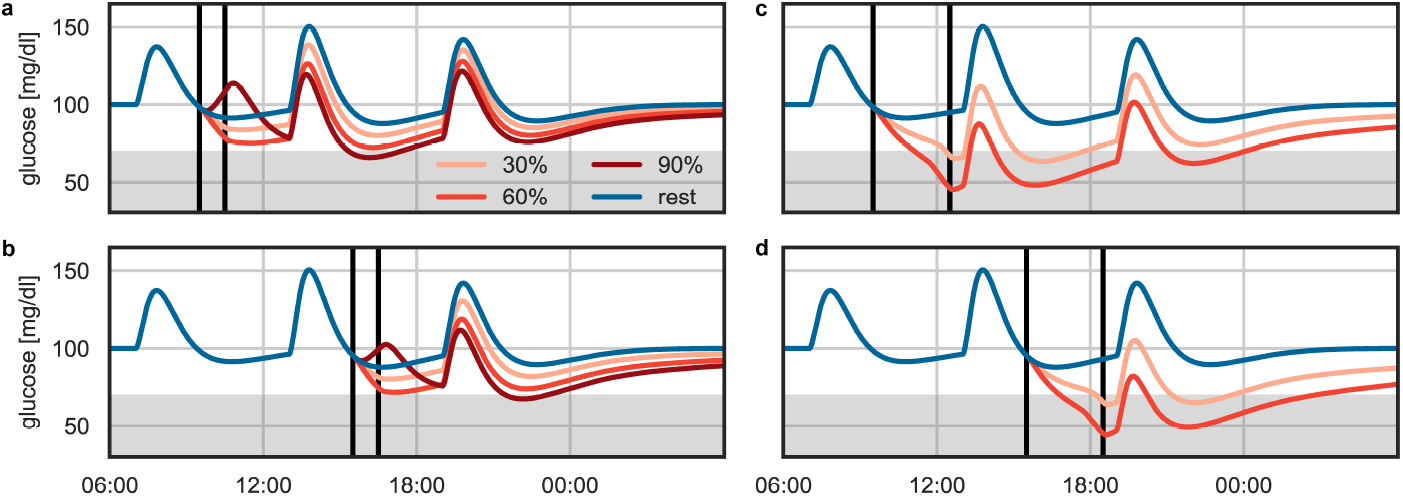
Full-day simulation of glucose dynamics at rest (blue), including three meals and corresponding insulin bolus injections. In addition, PA is performed (vertical lines) for (a, b) 60 min and (c, d) 180 min in the morning (a, c) and the afternoon (b, d). Glucose dynamics are shown for moderate intensities of 30% and 60% VO_2_^max^ and for high-intensity PA at 90% VO_2_^max^.

For moderate intensities, plasma glucose decreases during PA and slowly approaches resting glucose levels afterwards. For prolonged PA of 180 min, depletion shows slightly earlier for an intensity of 60% VO_2_^max^ than for 30% VO_2_^max^. Blood glucose decreases with increasing intensity and duration. Glucose levels are lower during the night for afternoon PA than for comparable morning PA, indicating higher risk for nocturnal hypoglycemia when exercise is performed later in the day.

Glucose concentration increases during high-intensity PA, but decreases to levels below those of moderate-intensity PA after the activity. The model thus confirms observations that high-intensity PA protects against hypoglycemia short-term [20], while the risk for late-onset hypoglycemia increases with higher intensity and duration of PA [24, 31]. Furthermore, risk of nocturnal hypoglycemia is increased following afternoon compared to morning PA [18].

### 3.5 Insulin Reduction for Post-Exercise Meal Bolus

Lastly, we studied reduction of the correction insulin bolus for a meal post-exercise, a strategy successful in protecting against early-onset but not late-onset hypoglycemia [10]. We simulated PA in the morning from 10:00 to 12:00 and an insulin bolus for lunch at 13:00 either given in full, or reduced by 25% or 50% (Fig. 7).

**Figure 7:**
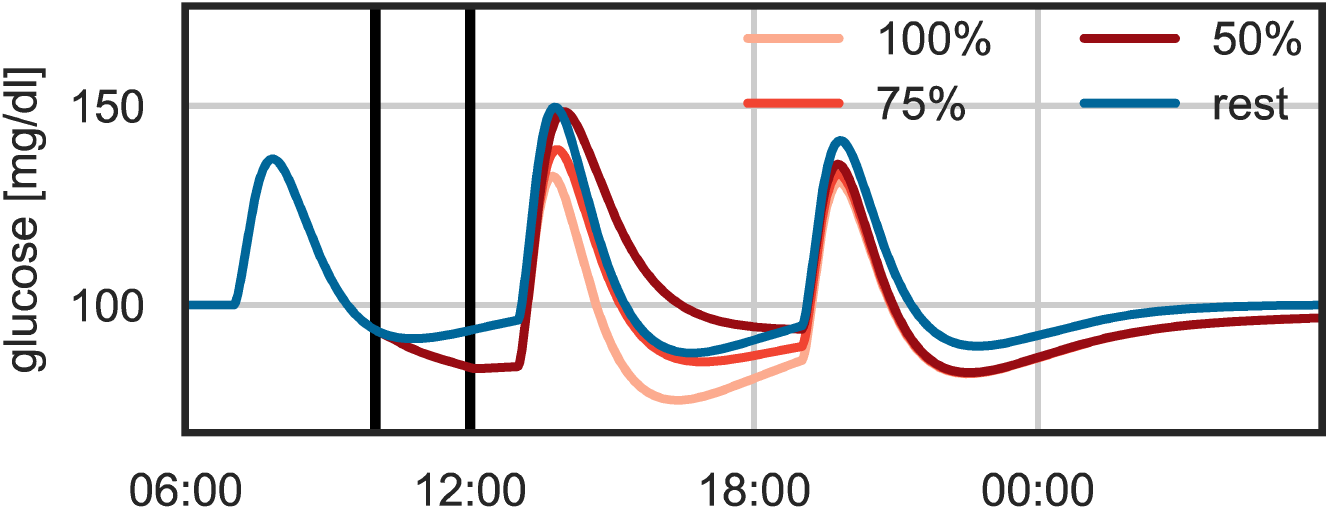
Full-day simulation with insulin reduction for post-PA meal bolus. Glucose dynamics are shown for rest (blue) and morning PA with a full (100%), 75% and 50% insulin dose administered for lunch.

As expected, plasma glucose is initially higher after lunch for reduced insulin compared to the full dose and is comparable to resting glucose levels. However, glucose concentrations are lower than resting levels throughout the night in all scenarios, independent of the bolus reduction. These results indicate that bolus reduction post-PA may protect against early-onset hypoglycemia, but does not target PA-induced late-onset hypoglycemia.

## 4 Discussion

We presented a model of exercise-related changes in glucose-insulin regulation in type 1 diabetes. The model comprehensively covers acute and prolonged PA-driven processes from low to high intensity to capture changes in glucose dynamics during and after PA. We employ transfer functions to keep the model compact while allowing transitions between different exercise regimes.

Our model quantitatively describes exercise-driven changes in insulin sensitivity, GU and GP for all exercise regimes and allows for varying intensity throughout the activity. It describes the prolonged rise in insulin sensitivity that drives glycogen repletion during recovery, a main cause of late-onset hypoglycemia. We further modeled the insulin-independent acute rise in GU during the activity and the simultaneous increase in hepatic GP responsible for meeting the increased energy demand of the exercising muscles. To capture glucose metabolism during prolonged PA, we added the reduction in GP due to liver glycogen depletion. We also included GP increases during high-intensity PA.

Our model accurately reflects glucose dynamics for low-to high-intensity PA during the activity and the following recovery period. We validated model predictions on independent data of moderate-intensity PA, with and without depletion. No additional data were available for independent validation of the high-intensity model extension.

The developed model includes simple modules for insulin bolus injections and meal intake, which makes it suitable for full-day simulations with simple meals.

To evaluate the long-term prediction capabilities of the model, we presented full-day simulations with different PA scenarios. For moderate-intensity PA, model predictions agree with the observations that glucose levels decrease with intensity and duration. In addition, we correctly predict that blood glucose is lower during the night following afternoon compared to morning PA, which is associated with a higher risk for nocturnal hypoglycemia. Lastly, the model successfully captures the blood glucose rise during high-intensity PA, and the subsequent decrease below levels reached with comparable moderate-intensity PA several hours after the activity.

Our determination of the model parameters suffers from two drawbacks: first, we only identify some of the parameters directly from data, but have to rely on domain knowledge and literature values for other parameters. We partially addressed this problem by testing model predictions on independent data sets, but cannot ensure that the individual model parts are physiologically accurate. Second, the available data sets only provide average glucose responses over several study participants. For future application of the model, it would be beneficial to use individual patient data for subject-specific model adjustment and to assess inter-patient variability. Based on these results, an in-silico patient population could be generated and applied for realistic simulation studies, in which the whole range of potential glucose outcomes would be covered.

## 5 Conclusion

Summarizing, our comprehensive model of glucose-insulin regulation captures the acute and prolonged effects of low-to high-intensity PA on glucose metabolism.

In addition, it includes meal intake and insulin injection kinetics, making it well-suited to describe glucose dynamics during everyday life. Hence, the model can be used to predict blood glucose during PA and recovery and to evaluate the impact of PA on glucose levels acutely and up to several hours after the activity in practical scenarios. It can further be used to compare different exercise types and the associated hypoglycemia risk within one framework. We anticipate that our model finds applications as an ‘exercise calculator’ for clinical decision support [13] to assist patients and clinicians achieve good glycemic control after exercise, as well as for improving control algorithms for closed-loop insulin delivery. Its modular nature makes it easily adaptable to other forms of insulin delivery, in particular insulin pumps, and readily allows extension of our comparatively simple meal model to more complex meals.

We evaluated the model’s performance on real data, but further validation of the model components on the individual patient level is warranted before application in a clinical setting.

## Acknowledgment

We thank Jörg Stelling, Gilbert Koch and Tamara van Donge for helpful feed-back and discussions.

## Notes

### Competing Interest Statement

The authors have declared no competing interest.

